# OXA β-lactamases from *Acinetobacter* spp. are membrane-bound and secreted into outer membrane vesicles

**DOI:** 10.1101/2024.11.04.622015

**Authors:** Lucia Capodimonte, Fernando Teixeira Pinto Meireles, Guillermo Bahr, Robert A. Bonomo, Matteo Dal Peraro, Carolina López, Alejandro J. Vila

## Abstract

β-lactamases from Gram-negative bacteria are generally regarded as soluble, periplasmic enzymes. NDMs have been exceptionally characterized as lipoproteins anchored to the outer membrane. A bioinformatics study on all sequenced β-lactamases was performed that revealed a predominance of putative lipidated enzymes in the class D OXAs. Namely, 60% of the OXA class D enzymes contain a lipobox sequence in their signal peptide, that is expected to trigger lipidation and membrane anchoring. This contrasts with β-lactamases from other classes, which are predicted to be mostly soluble proteins. Almost all (> 99%) putative lipidated OXAs are present in *Acinetobacter* spp. Importantly, we further demonstrate that OXA-23 and OXA-24/40 are lipidated, membrane-bound proteins in *Acinetobacter baumannii*. In contrast, OXA-48 (commonly produced by Enterobacterales) lacks a lipobox and is a soluble protein. Outer membrane vesicles (OMVs) from *Acinetobacter baumannii* cells expressing OXA-23 and OXA-24/40 contain these enzymes in their active form. Moreover, OXA-loaded OMVs were able to protect *A. baumannii, Escherichia coli* and *Pseudomonas aeruginosa* cells susceptible to piperacillin and imipenem. These results permit us to conclude that membrane binding is a bacterial host-specific phenomenon in OXA enzymes. These findings reveal that membrane-bound β-lactamases are more common than expected and support the hypothesis that OMVs loaded with lipidated β-lactamases are vehicles for antimicrobial resistance and its dissemination. This advantage could be crucial in polymicrobial infections, in which *Acinetobacter spp.* are usually involved, and underscore the relevance of identifying the cellular localization of lactamases to better understand their physiology and target them.

**IMPORTANCE:** β-lactamases represent the main mechanism of antimicrobial resistance in Gram-negative pathogens. Their catalytic function (cleaving β-lactam antibiotics) occurs in the bacterial periplasm, where they are commonly reported as soluble proteins. A bioinformatic analysis reveals a significant number of putative lipidated β-lactamases, expected to be attached to the outer bacterial membrane. Notably, 60% of class D OXA β-lactamases (all from *Acinetobacter* spp) are predicted as membrane-anchored proteins. We demonstrate that two clinically relevant carbapenemases, OXA-23 and OXA-24/40 are membrane-bound proteins in *A. baumannii*. This cellular localization favors secretion of these enzymes into outer membrane vesicles that transport them outside the boundaries of the cell. β-lactamase-loaded vesicles can protect populations of antibiotic-susceptible bacteria, enabling them to thrive in the presence of β-lactam antibiotics. The ubiquity of this phenomenon suggests that it may have influenced the dissemination of resistance mediated by *Acinetobacter* spp., particularly in polymicrobial infections, being a potent evolutionary advantage.

## INTRODUCTION

Bacteria possess a potent and diverse arsenal to resist the action of antibiotics. Gram-negative bacteria have predominantly evolved the expression of β-lactamases as one of the main mechanism of resistance against β-lactam antibiotics (1, 2). To date, more than 8200 different β-lactamase variants are reported (3), a number that is increasing at an alarming pace. The evolution and dissemination of genes coding for β-lactamases in opportunistic and pathogenic bacteria has been accelerated by the misuse and overuse of antibiotics worldwide, representing a major challenge for public health (1, 4).

β-lactamases are classified into four groups (A, B, C and D) according to the Ambler system, based on sequence homology (5). Class A, C and D enzymes are serine-β-lactamases (SBLs), which share a common protein fold, but display differences in their active sites and catalytic mechanisms (6, 7). In contrast, class B enzymes are metallo-β-lactamases (MBLs) requiring Zn(6) ions for their hydrolytic activity (8). From the clinical point of view, carbapenemases are the largest public health threat since they are able to inactivate carbapenems, the most potent β-lactams available in clinical practice (9, 10). Carbapenemases have been identified three out of these four classes, including all class B MBLs, members of class A (KPC, GES and NMC) and many class D β-lactamases (from the OXA family, named upon their oxacillinase activity) (11, 12).

In Gram-negative bacteria, mature β-lactamase enzymes are localized in the periplasmic space (13–16), where they cleave β-lactam antibiotics, thwarting their activity against enzymes involved in peptidoglycan cross-linking. β-lactamases have been historically regarded as soluble periplasmic enzymes. In contrast, a reduced number of β-lactamases have been exceptionally characterized as lipoproteins anchored to the inner leaflet of the outer membrane, such as the class A enzymes BRO-1 from *Moraxella catarrhalis* (17) and PenI from *Burkholderia pseudomallei* (18). More recently, the widespread Zn-dependent carbapenemase NDM (6) (with 68 reported clinical variants to date) was identified as a lipidated, membrane-bound enzyme, a localization that enhances the stability of this enzyme upon the zinc starvation process during an infection (19, 20).

β-lactamases are produced as cytoplasmic precursors with an N-terminal signal sequence (the signal peptide) that directs these precursors to one of the two main export pathways (Sec or Tat) responsible for protein translocation into the periplasmic space (13). Most biochemically characterized β-lactamases have been shown to translocate across the inner membrane via the Sec system (14, 16). In the case of soluble, non-lipidated β-lactamases, the type I signal peptidase (S cleaves the signal peptides of the precursor enzymes to generate the mature proteins (13, 21, 22). Instead, in the case of lipoproteins, the signal peptides are cleaved by the type II signal peptidase (SpII) during the lipoprotein maturation process (16, 22). Lipoprotein precursors possess a characteristic 4 amino acid motif known as a lipobox (23). The lipobox consensus sequence is [LVI]-[ASTVI]-[GAS]-C (24–26). The lipidation machinery in the periplasm transfers a diacylglycerol group to the free sulfhydryl of the cysteine residue in the lipobox and cleaves the signal sequence, leaving an acylated cysteine at the N-terminus. A further acyl group is then added generating a mature, triacylated lipoprotein that is inserted into the membrane (27). This lipidation mechanism has been thoroughly characterized in *E. coli* and it is widespread among Enterobacterales and non-fermenters (25, 28, 29).

In the case of NDM-1, membrane anchoring stabilizes this MBL against zinc deprivation and favors its incorporation into outer membrane vesicles (OMVs) (19). These nano-sized spherical structures bud and detach from the outer membrane of Gram-negative bacteria (30), playing several roles as decoys for phages and antibiotics, transporting nucleic acid, outer membrane and periplasmic proteins as well as insoluble cargo (31, 32). Similar processes have been also reported in Gram-positive organisms (33). NDM incorporation into vesicles increases the available enzyme levels at the infection site, extending antibiotic hydrolysis beyond the limits of the bacterial cell (20). As a result, NDM-loaded vesicles can protect populations of otherwise antibiotic-susceptible bacteria (19). Furthermore, vesicles can also mediate the transfer of the *bla*_NDM-1_ gene between bacteria (34, 35). The selection of the protein cargo in the case of NDM-1 depends on the interaction with the bacterial membrane, in which the covalent attachment through the lipid moiety is the main determinant (36, 37).

To explore the ubiquity of lipidation among β-lactamases, we performed a bioinformatic analysis of the signal peptides of all β-lactamases deposited in the β-lactamase database (www.BLDB.eu) (3). This study reveals a low number of lipobox sequences in enzymes from classes A, B, and C. In contrast, almost 60% of class D β-lactamases (all of them OXA enzymes) have a lipobox sequence and are thus predicted to be membrane-bound lipoproteins. Noteworthy, all putative OXA β-lactamase lipoproteins are found in *Acinetobacter* spp. Herein, we provide experimental evidence that the clinically relevant carbapenemases OXA-23 and OXA-24/40 from *A. baumannii* are membrane-bound lipoproteins. Disruption of the lipobox via mutagenesis results in the expression of soluble enzymes, supporting the bioinformatics analysis and molecular simulations.

The membrane-bound localization of OXA β-lactamases favors their incorporation into OMVs, as reported for NDM-1. The evolving hypothesis is that these OXA-loaded vesicles are capable of improving survival of not only *Acinetobacter* spp. in conditions of high β-lactam concentrations but also β-lactam-susceptible bacteria that are present in polymicrobial infections. We conclude that membrane-bound β-lactamases and their vesicle packaging represent a significant adaptation response in *Acinetobacter* spp. We propose that this cellular localization, linked to the secretion of lactamases into OMVs, represents an evolutionary advantage for *A. baumannii*, a highly troublesome pathogen associated with community-acquired and nosocomial infections. The protective effect of these vesicles confers a population advantage provided by *A. baumannii* OXA-producers in multiple clinical environments.

## RESULTS

### Bioinformatics predict that most class D β-lactamases (OXAs) are lipidated

To identify putative lipidated β-lactamases, we performed an *in silico* analysis of all β-lactamase sequences available in the BLDB database (3) using the SignalP 6.0 server (38). This server employs a machine-learning algorithm that classifies signal peptides into one of the five known types (Sec/SPI, Sec/SPII, Sec/SPIII, Tat/SPI and Tat/SPII) by means of a score that predicts the possibility of the signal peptide being transported and processed by each system. Our intention was to label all reported β-lactamases into soluble and lipidated enzymes. Despite the focus of the current work being on β-lactamases in Gram-negative organisms, the analysis covered all sequenced enzymes.

From a total number of 7479 accessible β-lactamase sequences (April 2024), 7226 were predicted to contain an N-terminal signal peptide targeting the Sec translocation pathway (97 %) (Fig. 1A), confirming that most β-lactamases are Sec substrates (16). The small proportion of β-lactamases predicted to be translocated by the Tat system mostly belong to highly divergent class A enzymes, suggesting that this feature does not imply any evolutionary connection among them but, instead, an adaptation to each organism. Indeed, there are enzymes from Gram-negative bacteria such as *Burkholderia* spp. (PenA and PenI enzymes), *Xanthomonas* spp. (XCC enzymes) and *Mycobacterium* spp. (MFO from M. fortuitum and MAB from *M. abscessus*) and from the Gram-positive organism *Streptomyces* spp. (SDA enzymes) (Supplementary Table 1).

**Fig 1.**
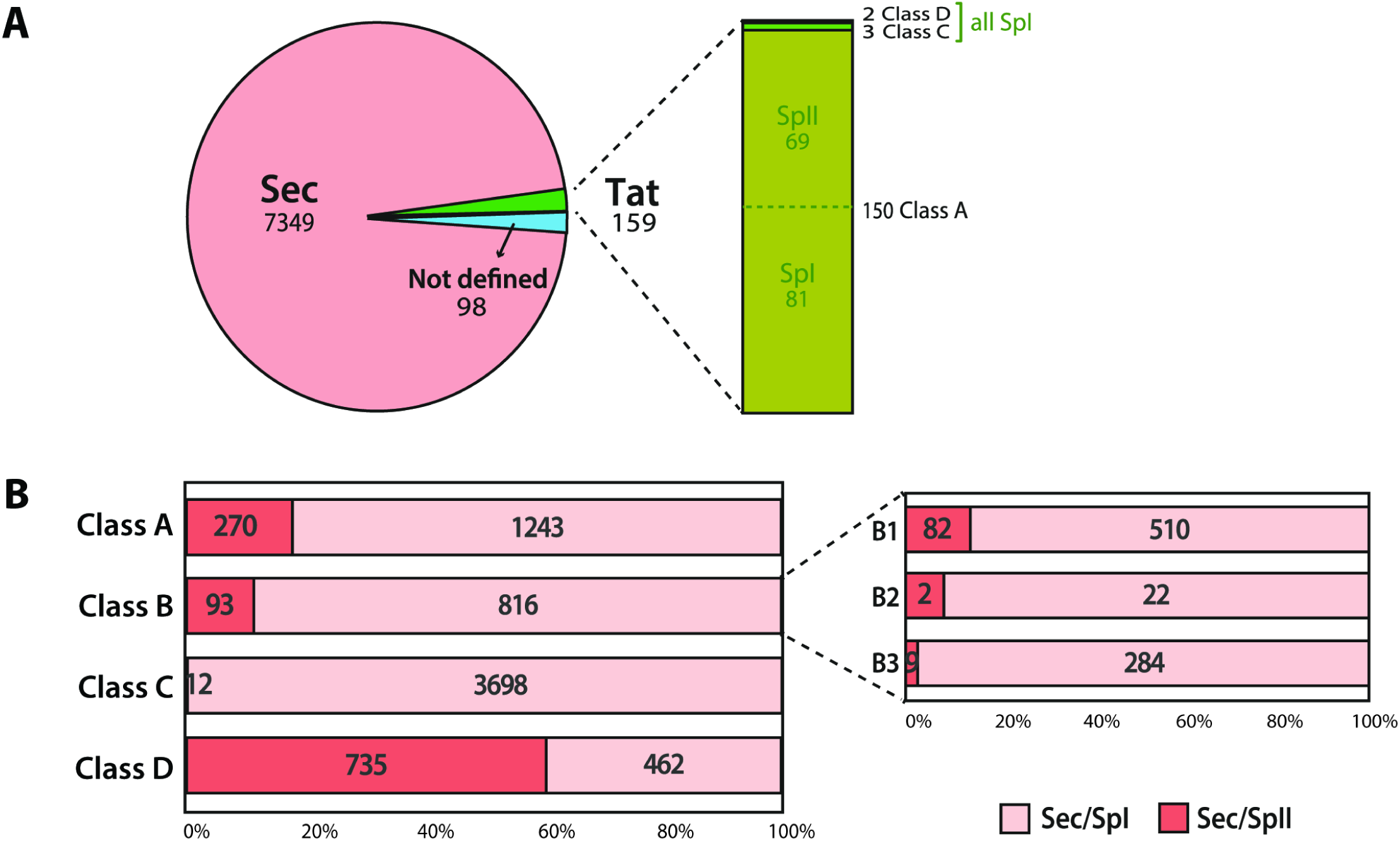
Class D contains the largest family of putative lipidated β-lactamases. **(A)** Pie chart indicating the number of β-lactamase sequences predicted to be translocated by the Sec and Tat systems. The bar at the right shows the distribution of β-lactamases translocated by the Tat system in each class, detailing if they are putative substrates of SpI (putative soluble proteins) or SpII (putative lipoproteins). **(B)** Putative substrates of SpI or SpII translocated by the Sec system, separated according to the β-lactamase class. The absolute numbers of each category are indicated inside each bar. The three subclasses of class B enzymes (B1, B2 and B3) are indicated at the right. Class D enzymes show the largest number of predicted lipoproteins.

The identification of a lipobox sequence helps annotate proteins as substrates of the signal peptidase II, therefore targets for lipidation and membrane anchoring. Lipoboxes were identified in the sequences of 17% of class A enzymes, 10 % of class B, and only two (0.3%) class C β-lactamases (Fig. 1B and Table 1). This result agrees with the consensus that considers β-lactamases as soluble periplasmic proteins. In stark contrast, ca. 60% of class D enzymes are putative lipoproteins (Fig.1B and Table 2). Table 1 shows several representative class A and B β-lactamases predicted as lipidated proteins, while Table 2 lists all families of putative lipidated OXAs. We also indicate the most likely translocation and processing pathway in each case. Supplementary Table 1 includes the predictions for all β-lactamases, with the scores provided by the SignalP 6.0 server.

**Table 1.** β-lactamases from classes A and B with lipoboxes in their signal peptides. The asterisk next to the lactamase names indicate the enzymes for which the membrane localization has been experimentally assessed. The most frequent organisms expressing the lactamase gene are indicated in the third column.

| Ambler class or subclass | β-lactamase | Organism | Translocation system |
| --- | --- | --- | --- |
| A | BcIII* | *Bacillus cereus* | Sec/SpII |
| A | BlaC | *Mycobacterium tuberculosis* | Tat/SpII |
| A | BlaP-1/2 | *Bacillus licheniformis* | Sec/SpII |
| A | BRO-1*/2* | *Moraxella catarrhalis* | Tat/SpII |
| A | PC-1* | *Staphylococcus aureus* | Sec/SpII |
| A | PenA-1/39 | *Burkholderia multivorans* | Tat/SpII |
| A | PenI-1/8* | *Burkholderia pseudomallei* | Tat/SpII |
| A | ROB-1/5, 8/13 | *Pasteurellales, Moraxella spp.* | Sec/SpII |
| B1 | AFM-1/4 | *Burkholderiales, Pseudomonas spp.* | Sec/SpII |
| B1 | NDM-1/68* | Enterobacterales, non-fermenters | Sec/SpII |
| B3 | ECM-1 | *Erythrobacter citreus* | Sec/SpII |
| B3 | EFM-1 | *Erythrobacter flavus* | Sec/SpII |
| B3 | EVM-1 | *Erythrobacter vulgaris* | Sec/SpII |

**Table 2.** Predicted cellular localization of principal class D β-lactamases. In the organism column are specified the most reported organism carrying the lactamase gene or the order they belong to. The three OXA proteins selected in this study are highlighted in red (OXA-23, 24/40 and 48, the representative members of each of these families). Supplementary Table 2 describes all OXA-subfamilies.

| Class D β-lactamase | Number of enzymes | Organism(s) | Predicted cell localization |
| --- | --- | --- | --- |
| OXA-1-like | 12 | *Pseudomonadales, Shewanellaceae, Pasteurellales, Enterobacterales* | soluble |
| OXA-2-like | 31 | *Pseudomonadales, Burkholderiales, Shewanellaceae, Enterobacterales, Vibrionales, Pasteurellales* | soluble |
| OXA-10-like | 60 | *Burkholderiales, Pseudomonadales, Shewanellaceae, Enterobacterales* | soluble |
| OXA-22-like | 7 | *Ralstonia* spp. | soluble |
| OXA-23-like | 49 | *Acinetobacter* spp. * | lipidated |
| OXA-24/40-like | 24 | *Acinetobacter* spp. | lipidated |
| OXA-48-like | 66 | *Enterobacterales, Shewanellaceae* | soluble |
| OXA-50-like | 60 | *Pseudomonas* spp. | soluble |
| OXA-51-like | 382 | *Acinetobacter* spp. | lipidated |
| OXA-58-like | 8 | *Acinetobacter* spp. | lipidated |
| OXA-60-like | 7 | *Ralstonia* spp. | soluble |
| OXA-134-like | 34 | *Acinetobacter* spp. | lipidated |
| OXA-61-like | 49 | *Campylobacter* spp. | soluble |
| OXA-211-like | 17 | *Acinetobacter* spp. | lipidated |
| OXA-213-like | 51 | *Acinetobacter* spp. | lipidated |

The putative lipoproteins belonging to class A include an important number of enzymes from Gram-positive and Gram-negative bacteria, a few of them already characterized experimentally. BcIII from *Bacillus cereus* was the first characterized membrane-bound lactamase (39). Later, BRO-1 was the first β-lactamase from a Gram-negative bacteria characterized as a membrane-bound protein (17). BRO-1 and BRO-2 from *Moraxella catarrhalis*, are both predicted as lipoproteins dependent on the Tat system. PenI enzyme from *B. pseudomallei* was experimentally shown to be a membrane-bound protein (18). Here we show that the variants of PenA, PenB, PenI and other members of the Pen family produced by *Burkholderia* species are predicted as lipoproteins translocated by the Tat system.

In the case of class B MBLs, subclass B1 includes all variants of AFM and NDM enzymes, CHM-1, and ZOG-1, as putative lipoproteins dependent on the Sec system. No proteins from subclass B2 were predicted to be lipidated with a high score. However, within subclass B3, certain enzymes were identified as lipoproteins (Table 1).

Most class D β-lactamases are also known as oxacillinase enzymes or OXAs (12). Remarkably, all putative lipidated OXAs are chromosomally-encoded or acquired from *Acinetobacter* species, except for OXA-63-like enzymes from *Brachyspira pilosicoli*, OXA-347, −1089 and −1090 (26 out of 735 OXA enzymes). Indeed, the main groups of carbapenem-hydrolyzing class D β-lactamases (CHDL) were predicted as lipoproteins: the chromosomally-encoded OXA-51-like and the acquired OXA-23-like, OXA-58-like and OXA-24/40-like (Table 2). In contrast, soluble OXA enzymes are not predicted in *Acinetobacter* species. This direct link between protein lipidation and a bacterial host (*Acinetobacter*, in this case) is unique to class D enzymes, since lipidated class A and B enzymes are found in a wide variety of bacterial hosts (Table 1 and Table 2).

### Molecular simulations reveal a mechanism for membrane association for lipidated OXA β-lactamases

To better characterize their possible association with membranes, we selected and performed coarse-grained molecular dynamics (CG-MD) simulations of the two most clinically important β-lactamases related to carbapenem resistance: OXA-23 and OXA-24/40. We ran the simulations with a lipid bilayer mimicking the composition of the inner leaflet of the *A. baumannii* outer membrane (12% cardiolipin, 16% phosphatidylglycerol (PG), and 72% phosphatidylethanolamine (PE) (see Methods)). The enzymes were triacylated at their N-terminal cysteine residue, as suggested by the bioinformatic analysis of the signal peptide. In all simulations (5 MD replicas) both enzymes readily anchored into and kept attached to the lipid bilayer via the triacyl moiety in their N-terminus. OXA-23 and OXA-24/40 adopted similar orientations on the membrane, laying their globular domain on its surface while leaving their active site facing the periplasmic space (Supplementary Movies 1 and 2; Fig. 2).

**Fig 2.**
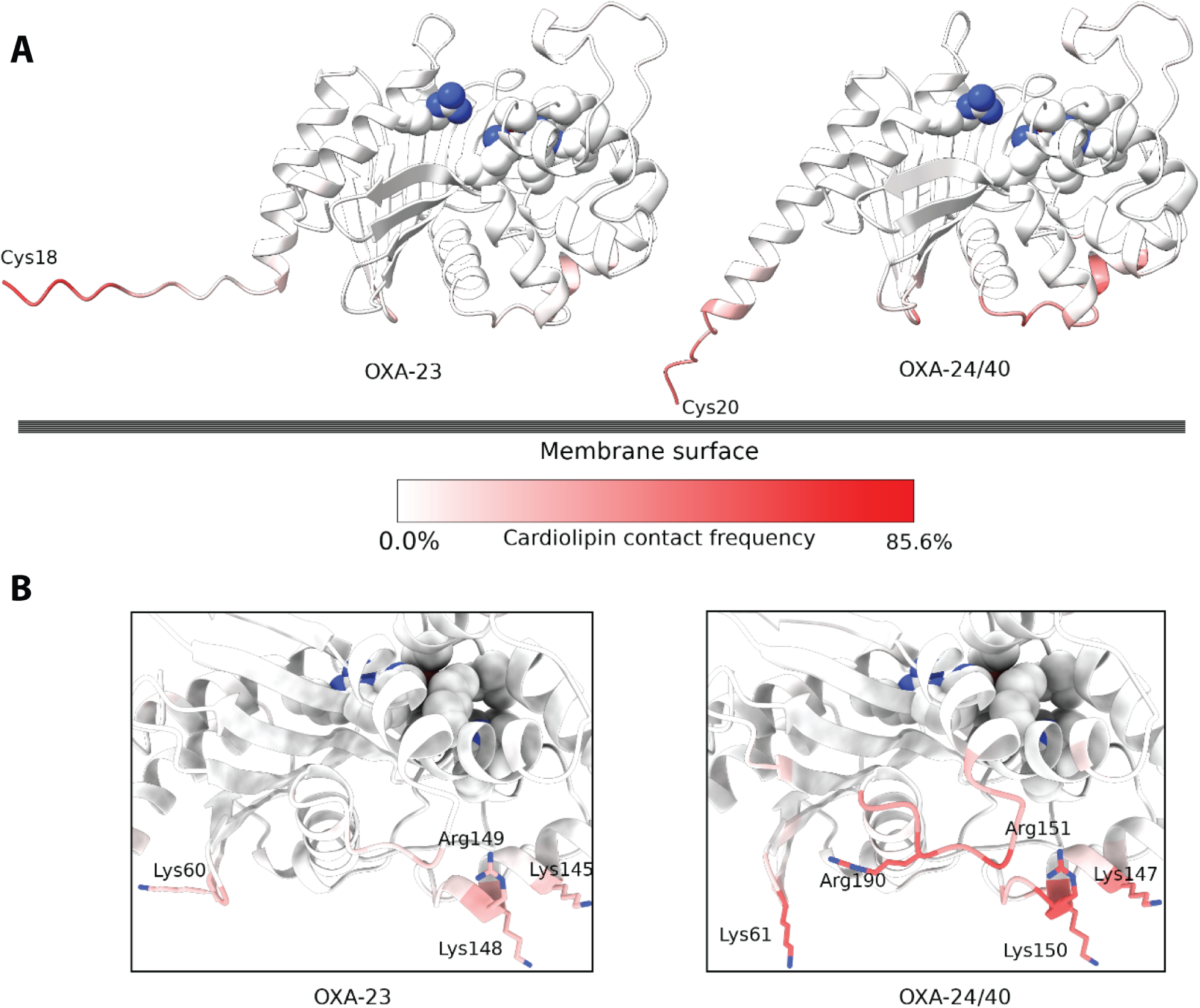
Residue-wise contact frequency (%) with cardiolipin lipids during the simulations. (A) OXA-23 and OXA-24/40 contact frequency (measured as percentage of simulation time) with cardiolipin, color coded from white (0% frequency) to red (85.5% frequency). The lipidated cysteines (Cys18 and Cys20) are indicated in the N-terminus of both β-lactamases. Active site residues are shown as spheres. (B) A close-up of the residues (shown as sticks) on the opposite side of the active-site (shown as spheres). These positively charged residues also contribute to membrane binding, positioning the active-site towards the periplasm and preventing its occlusion.

This positioning of OXA-23 and OXA-24/40 might be a structural feature that prevents the occlusion of their active-site by interaction with the lipid bilayer, as the same phenomenon has been observed in MD studies of lipidated NDM-1 (36).

Cardiolipin molecules displayed considerably high contact frequency (percentage of the simulation time) with the globular domain of OXA-24/40, despite representing only 12% of the membrane’s lipid composition, notably with residues Arg151 (56.9%), Arg190 (53.8%), Lys61 (53.2%), Lys150 (45%), and Lys147(43.3%) (Fig 2B). Albeit to a lesser extent, the residues in the analogous positions in OXA-23 also helped stabilize the enzyme’s conformation on the membrane surface via cardiolipin interactions as measured by contact frequency: Arg149 (26.9%), Lys148 (25%), Lys145 (21.9%), and Lys60 (20.1%) (Fig 2B). Despite not being essential for membrane anchoring, these electrostatic interactions could help orient OXA-23 and OXA-24/40 in such a way that their active-sites remain available to substrate binding.

### OXA-23 and OXA-24/40 are lipidated, membrane anchored proteins

To experimentally test the bioinformatic predictions and the molecular simulations, we performed cellular fractionation experiments. We selected the two putative lipidated OXA-23 and OXA-24/40 expressed in *A. baumannii*, and a putative soluble periplasmic OXA, OXA-48, lacking a lipobox sequence and commonly produced by Enterobacterales such as *K. pneumoniae* and *E. coli* (Fig. 3A) (12). All these selected proteins are clinically relevant carbapenemases (1, 9).

**Fig 3.**
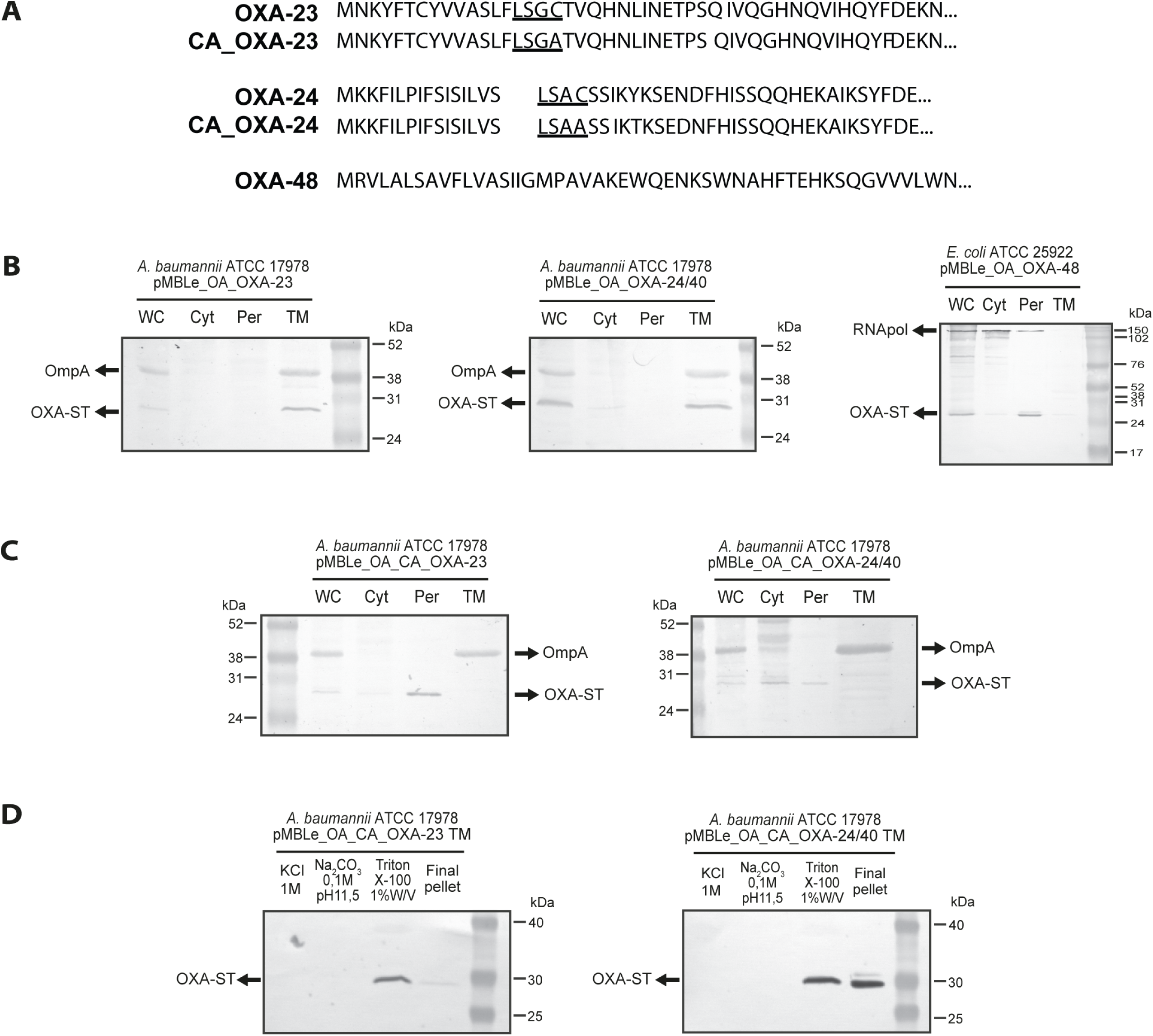
OXA-23 and OXA-24/40 are membrane anchored proteins while OXA-48 is soluble periplasmic. (A) N-terminal sequences of the OXAs object of the experimental analysis. The lipoboxes are underlined and the cysteine target of lipidation is bolded. (B) Cell fractionation of *A. baumannii* ATCC 17978 expressing OXA-23 and OXA-24/40, and *E. coli* ATCC 25922 expressing OXA-48 (C) Cell fractionation of *A. baumannii* expressing the Cys18Ala-OXA-23 (CA_OXA-23) and Cys20Ala-OXA-24/40 (CA_OXA-24/40) variants. (D) Solubilization assays of OXA-23 and OXA-24/40 from *A. baumannii* total membranes. OXA-ST indicates the bands corresponding to anti-ST antibodies, which match with the molecular weight of the enzymes fused to the tag. OmpA indicated the bands revealed by anti-OmpA antibodies which recognize the outer membrane protein OmpA. RNApol indicated the bands revealed using RNA polymerase antibodies.

These three proteins were expressed by the pMBLe_OA plasmid with their native signal peptides and with a Strep-tag (ST) fused to the C-terminal to allow uniform immunodetection (37). We chose *A. baumannii* ATCC 17978 and *Escherichia coli* ATCC 25922 as laboratory strains for the different organisms with isogenic backgrounds. The grown cells were subjected to a cell fractionation assay, which allowed collecting the periplasmic fraction (Per) after treatment with lysozyme. Sonication then led to the separation of total membranes (40) from the cytoplasm (Cyt). Whole cells (WC) and each fraction were analyzed by immunoblotting with anti-ST antibodies for the β-lactamases. Specific antibodies against cytoplasmic RNA polymerase (RNApol) and the outer membrane protein A (OmpA), were used as controls for the quality of the preparation of the soluble and membrane fractions, respectively. Fig. 3B shows that OXA-23 and OXA-24/40 were present in the membrane fractions of *A. baumannii*, with no accumulation of these proteins in the periplasmic fraction. In contrast, OXA-48 was detected only in the periplasmic fraction of *E. coli* as a soluble β-lactamase (Fig. 3B).

The lipobox is a signature sequence of bacterial lipoproteins, and the cysteine residue located at its C-terminus is the target of lipidation. To confirm the role of this residue in the localization in the membrane fraction of OXA 23 and OXA-24/40 in *A. baumannii*, we substituted the Cys for Ala in both proteins (Cys18 in OXA-23 and Cys20 in OXA-24/40) (Fig. 3A) and analyzed the impact of this replacement in the cellular localization of both enzymes. Expression of the Cys18Ala_OXA-23 (CA_OXA-23) and Cys20Ala_OXA-24/40 (CA_OXA-24/40) variants in *A. baumannii* resulted in the accumulation of both proteins only in the periplasmic fractions as soluble proteins, separately from OmpA, which is present in total membrane fractions, confirming our hypothesis (Fig. 3C).

To assess the nature of the interaction between the bacterial membrane and OXA-23 and OXA-24/40, we attempted to solubilize these enzymes from the pure *A. baumannii* membranes using different methods. Treatment with high ionic strength (1 M NaCl) and highly basic pH (0.1 M Na_2_CO_3_ pH 11.5) did not release any of the two OXAs from the membrane fractions (Fig. 3D), indicating that they are not peripheral proteins associated with the membrane by only means of electrostatic interactions, although these interactions can better expose the active side to the periplasmic space, as observed with MD simulations (Fig. 2). Instead, OXA-23 and OXA-24/40 were only solubilized upon treatment with 1% w/v Triton X-100 (Fig. 3D), confirming that both enzymes interact with the membrane through hydrophobic interactions. Overall, these results establish that OXA-23 and OXA-24/40 are lipidated, membrane-bound proteins in *A. baumannii*.

The MIC values of piperacillin and imipenem against *A. baumannii* cells expressing the soluble and membrane-bound variants of both OXA-23 and OXA-24/40 were similar (Suppl. Table 3), revealing that the cell localization does not contribute to the resistance phenotype.

### Lipidated OXAs are selectively secreted into OMVs

We then explored whether membrane anchoring of OXAs results in packaging into OMVs. We purified OMVs from *A. baumannii* expressing native OXA-23 and OXA-24/40, and the soluble variants CA_OXA-23 and CA_OXA-24/40 and quantified the protein levels in the vesicles. Despite the protein levels of lipidated and soluble variants were comparable in whole cells, only the lipidated OXAs were incorporated at high levels in OMVs from *A. baumannii* (Fig. 4A). Fig. 4B shows that removal of the lipidation site for CA_OXA-23 leads to a substantial decrease of approximately 95% in the transported enzyme by vesicles. Similarly, in the case of CA_OXA-24/40, there is an approximately 85% reduction in the level of the enzyme into OMVs, indicating that membrane anchoring contributes to the selective secretion of OXA-23 and OXA-24/40 in *A. baumannii* cells.

**Fig 4.**
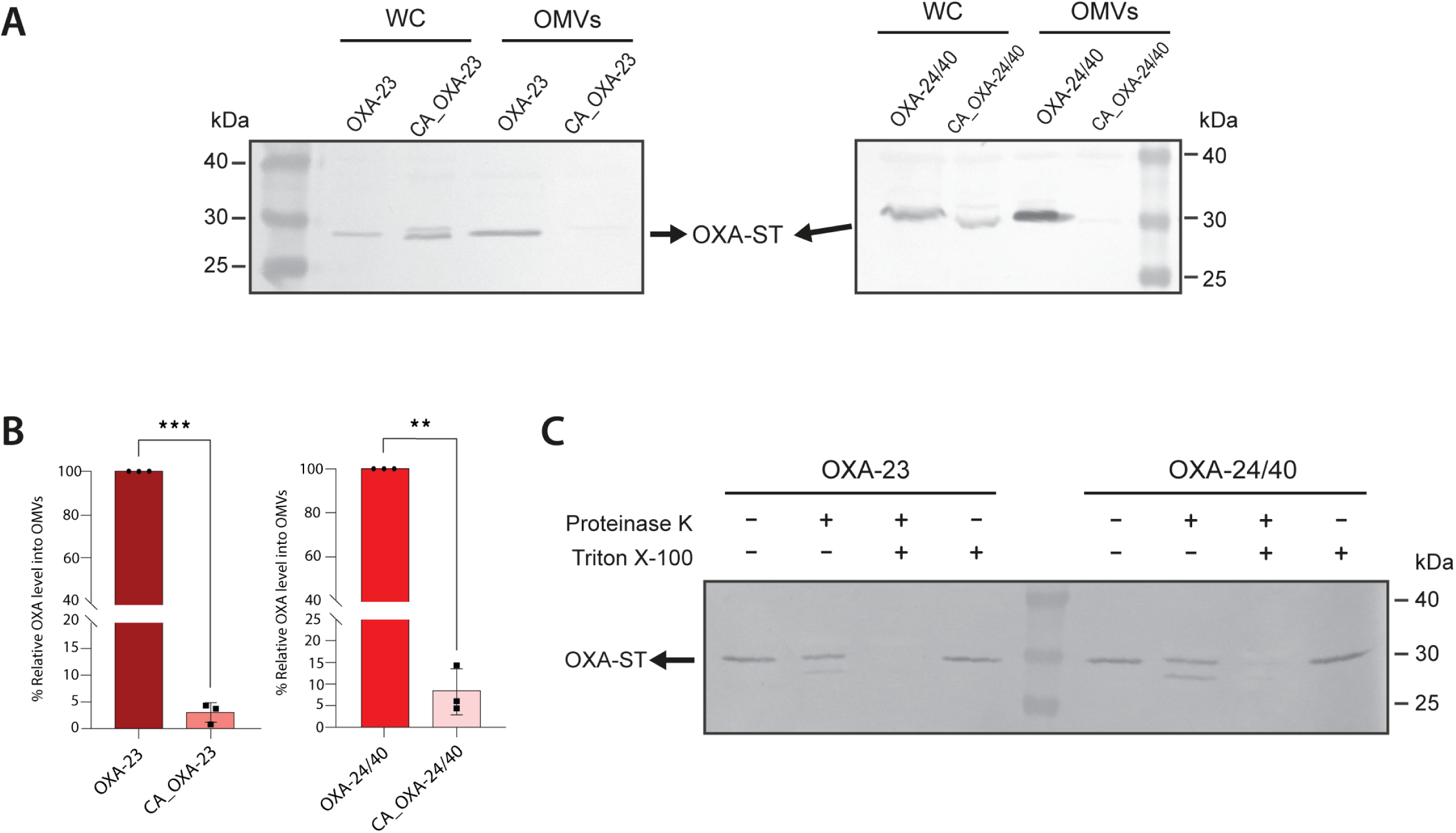
Membrane anchored OXAs are incorporated in higher proportions into OMVs than periplasmic soluble OXAs. (A) Anti-ST immunoblotting of whole cells (WC) and outer membrane vesicles (OMVs) from *A. baumannii* ATCC 17978 expressing OXA-23 or OXA-24/40 and its soluble variants (CA_OXA-23 or CA_OXA-24/40). (B) Comparison between the percentages (%) of the levels of the soluble variants CA_OXA-23 and CA_OXA-24/40 into OMVs. The plotted values, normalized to the corresponding wild-type OXA (lipidated OXA) levels, were obtained as described in Materials and Methods. Data correspond to three independent experiments (black filled symbols) and are shown as the mean value. Error bars represent standard deviations (SD). P-values according to the Student’s t-test: **p ≤ 0.01, ***p ≤ 0.001. (C) Anti-ST immunoblotting of OMVs from *A. baumannii* carrying OXA-23 or OXA-24/40 treated with and without Proteinase K and 1% v/v Triton X-100.

To assess whether the OXA enzymes are located in the lumen of the vesicles or pointing outwards, we exposed the OXA-loaded vesicles to proteinase K to assess the accessibility of this protease to the enzymes in the OMVs. Treatment of the intact vesicles with proteinase K did not alter the levels of OXA-23 or OXA-24/40. Instead, both enzymes were completely degraded by proteinase K in Triton-lysed vesicles (Fig. 4C). These experiments suggest that the proteins are located in the lumen of the OMVs, confirming the protective role of the vesicles from extracellular degradation when these enzymes are secreted.

### OXA-loaded OMVs protect β-lactam-susceptible bacteria

We then attempted to determine if the OXAs incorporated into vesicles were in their active forms by examining the protective effect of β-lactamase-loaded vesicles in β-lactam-susceptible bacteria. We tested this in β-lactam-susceptible *A. baumannii, E. coli,* and *P. aeruginosa* cells treated with OMVs from *A. baumannii* carrying the empty vector (EV) or expressing OXAs in the presence of imipenem or piperacillin, and we determined the MICs.

Incubation of β-lactam-susceptible *A. baumannii* with OMVs loaded with lipidated OXA-24/40 resulted in a MIC against imipenem of 32 μg/ml versus a MIC of 2 μg/ml for those cells incubated with vesicles loaded with the soluble variant, and a MIC of 0.125 μg/ml for control vesicles isolated from *A. baumannii* cells transformed with the empty plasmid (Fig. 5A and Supplementary Table 4). The same trend was observed for β-lactam-susceptible *E. coli* and *P. aeruginosa* cells and for OXA-23 and its soluble variant. In this case, the β-lactam-susceptible *A. baumannii* cells grew at 8 μg/ml imipenem when incubated with OXA-23-containing OMVs versus 1 μg/ml imipenem for vesicles loaded with the soluble variant CA_OXA-23 (Fig. 5A and Supplementary Table 4). A similar protection effect was observed for piperacillin, with a high impact in the MICs for the three tested β-lactam-susceptible bacteria when were incubated with OMVs loaded with lipidated OXAs (Fig. 5B and Supplementary Table 4). The trend of enhanced protection with a higher MIC when using OXA-24/40-loaded vesicles compared to OXA-23-loaded vesicles is likely attributed to the higher levels of OXA-24/40 in the concentration of OMVs used during the incubation experiment with susceptible bacteria. In fact, the levels of OXA-24/40 in the OMVs were approximately four times higher than those of OXA-23. Overall, these results indicate that OXA enzymes are active within vesicles produced by *A. baumannii,* playing a collective role improving the viability of antibiotic-susceptible bacteria and that this role is highly dependent on the membrane localization of these enzymes.

**Fig 5.**
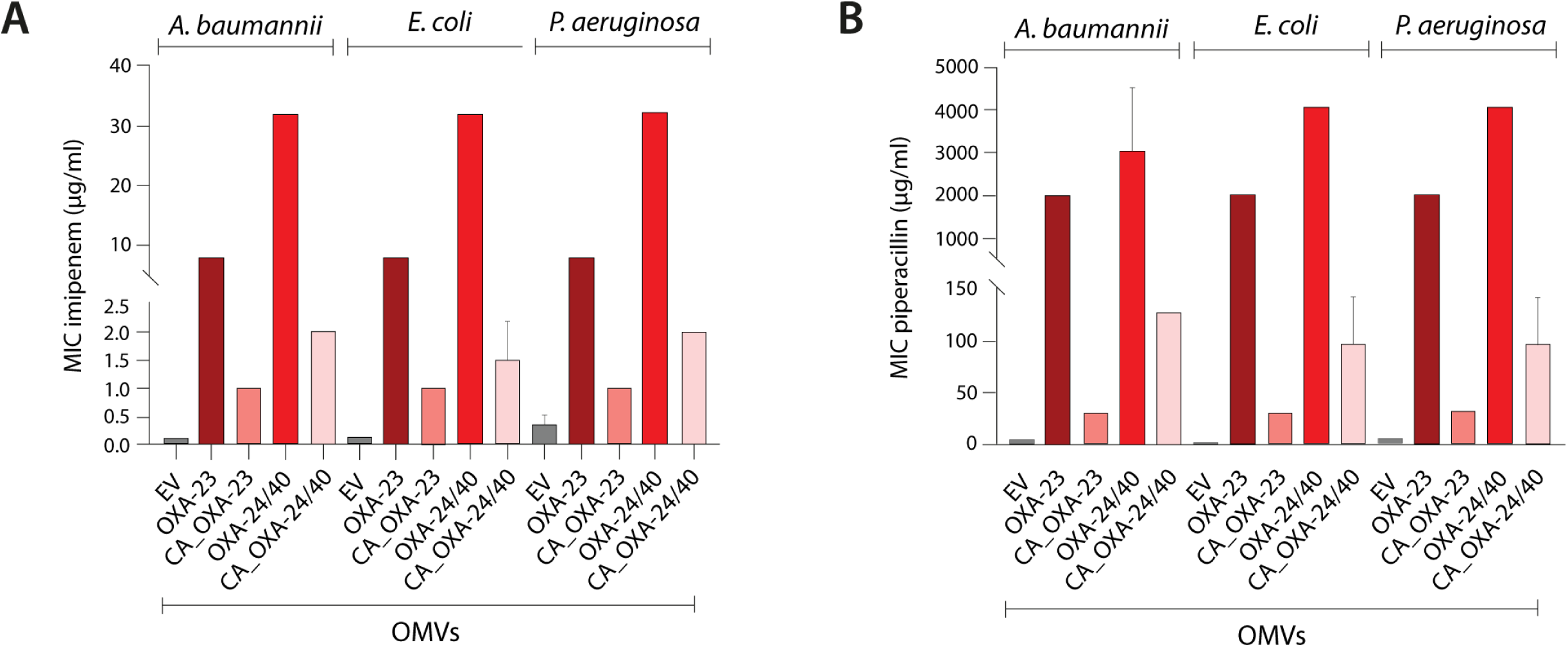
OMVs loaded with lipidated OXAs provide enhanced protection to β-lactam-susceptible bacteria than OMVs containing soluble variants. MIC values (µg/ml) against (A) imipenem and (B) piperacillin of β-lactam susceptible *A. baumannii*, *E. coli* and *P. aeruginosa* cells after treatment with OMVs purified from *A. baumannii* carrying empty the vector (EV) or expressing lipidated enzymes: OXA-23 or OXA-24/40 or soluble enzymes: CA_OXA-23 or CA_OXA-24/40. Data correspond to mean values from two independent experiments. Error bars represent standard deviations (SD). The MICs of susceptible bacteria against imipenem or piperacillin (without incubation with OMVs) are shown in Supplementary Table 3.

## DISCUSSION

Antimicrobial resistance (AMR) is an inevitable consequence of the use of antibiotics. β-lactamases represent the predominant mechanism of resistance to β-lactam antibiotics, as accounted by the early report of a penicillinase in 1940 before the clinical use of antibiotics (41). In the current century, we have witnessed an alarming spread of resistance genes coding for β-lactamases among pathogenic and opportunistic bacteria, with > 8,000 variants reported so far. Biochemistry and structural studies have described the mechanistic and structural features eliciting resistance to new drugs (42). However, there are still many aspects incompletely understood about the biochemical and mechanistic aspects of β-lactamases. Indeed, the catalytic mechanism of metallo-β-lactamases was dissected only 5 years ago(43), and a recent paper disclosed the role of anions in the biphasic nature of the kinetics of OXA enzymes(44, 45).

In addition to these biochemical issues, host-specific aspects of β-lactamases are poorly understood. For example, it is not clear why some enzymes, despite being plasmid-borne, are confined to some bacterial host(s), while others are present in a wider variety of microorganisms. This phenomenon has been recently addressed in the case of metallo-β-lactamases (46), but it is not defined for serine-dependent enzymes. Finally, the bacterial physiology of β-lactamases in the periplasm of Gram-negative bacteria is scarcely characterized, a problem that includes the incomplete knowledge of the cellular localization of these enzymes, which can be soluble or membrane-bound. The latter was shown to be the case of NDM-1 and its different allelic variants (47). The cellular localization of NDM variants improves the stability of these enzymes and favors their incorporation into OMVs from different bacteria (19, 46).

The identification of a lipobox sequence helps annotate a protein as putative lipidated and membrane-bound. In this work we examined the distribution of lipoboxes in the signal peptides of β-lactamases from all 4 classes. We found that the distribution of putative lipidated enzymes is diverse, depending on the class of β-lactamase and the microorganism. Within the class A group, representing one fourth of reported β-lactamases, only 15% of enzymes contain lipoboxes. Two of them, BRO from *Moraxella catarrhalis* and PenI from *B. pseudomallei*, have already been characterized biochemically. Within class B, all NDM variants are membrane-bound (68 out of > 800 enzymes), which have been reported in Enterobacterales and non-fermenters. Instead, class D enzymes are singular since 60% of the OXA β-lactamases contain a lipobox, all of them (with only one exception) being expressed in *Acinetobacter* spp. This suggests a host-specific effect for these enzymes that contrast with the host distribution of lipidated class A and class B β-lactamases.

Coarse-grained MD simulations predict that the lipid moiety attached to a Cys residue in the lipobox of OXA-23 and OXA-24/40 is inserted in the lipid bilayer in such a way that a patch in the globular domains of these enzymes presents an attractive interaction with the bacterial membrane. This interaction is expected to favor and stabilize membrane binding as well as to orient the enzyme active site towards the periplasm.

We validated these predictions by cellular fractionation and immunoblotting experiments, that reveal that OXA-23 and OXA-24/40 are lipidated, membrane-bound proteins in *Acinetobacter* spp. Mutagenesis of the Cys residue in the lipobox gives rise to soluble proteins in both cases. We also studied as a control OXA-48 (lacking a lipobox), a soluble, periplasmic enzyme in *E. coli*.

Both OXA-23 and OXA-24/40 were secreted into OMVs from *A. baumannii*. In both cases, the soluble variants resulting from a mutation at the lipobox were also secreted into OMVs, but to a much smaller content. This behavior is similar to what has been reported for NDM, i.e., membrane anchoring significantly facilitates secretion into vesicles, but the soluble variants can also be secreted (19). Incubation of different antibiotic-susceptible bacteria with OXA-loaded OMVs resulted in a protective effect of these bacteria against a carbapenem and a penicillin. In all cases, the protective effect (measured as an increase in the MIC) correlates with the amount of protein present in the OMVs. This reveals that both OXA-23 and OXA-24/40 are secreted in their active form, as is the case for NDM-1.

The current results help unify many findings in the literature and coalesce different observations. OXA-23 has been shown to display an extensive interaction network in *A. baumannii* by cross-linking and mass spectrometry experiments (48). OXA-23 interacts with the outer membrane proteins OmpA, OmpW, CarO and ABUW_2898, as well as with YiaD, an outer membrane protein that has been related to carbapenem resistance. In all these cases, the cross-linking was observed through residue Lys60 in OXA-23, which is one of the positively charged residues in the protein surface identified as making transient interactions with the outer membrane (Fig. 2). We propose that the reported interaction (48) is due to the proximity of OXA-23 to the outer membrane.

A proteomics analysis of OMVs from the multi-drug resistant clinical isolate *A. baumannii* DU202 revealed that OXA-23 accounted for 36% of the total protein content in OMVs (49). Thus, despite the relatively low levels of expression of OXA-23 in the current model organism, secretion of OXA β-lactamases into OMVs in clinical strains can be a relevant phenomenon. Liao and coworkers (50) informed the finding of OXA-58 in the lumen of OMVs from *A. baumannii* Ab290. Our analysis (Table 2) predicts the OXA-58 family as lipidated, also supporting this finding.

OMVs have been suggested to play different roles in bacteria. These vesicles are essential in cellular detoxification processes, by removing toxic periplasmic components that elicit envelope stress and compromise bacterial fitness. This has been shown to be the case for some class B β-lactamases when expressed in non-frequent bacterial hosts, such as VIM-2 and SPM-1 (46). Colquhoun *et al*. recently observed that hyperexpression of OXA-23 in *A. baumannii* induces collateral physiological damages by altering the peptidoglycan integrity (51). We postulate that the incorporation of OXA-23 into OMVs might mitigate the negative impact of OXA-23 overexpression by reducing the effective concentration of OXA-23 in the periplasm, at the same time increasing the levels of this β-lactamase in the environment by its presence in OMVs, thus contributing to resistance.

We also show here that OXA-loaded vesicles are able to protect bacterial populations of otherwise susceptible *A. baumannii*, *P. aeruginosa* and *E. coli* from the activity of β-lactam antibiotics. Therefore, the expression of lipidated OXA enzymes may provide an advantage both for the producing organism and in polymicrobial infections. Indeed, multi-drug resistant *Acinetobacter* spp. is increasingly reported in co-colonization events in intensive care units with Enterobacterales expressing extended spectrum β-lactamases (ESBLs) (52–54). A recent analysis from Semenec and coworkers (55) has described the relevance of cross-protection of *A. baumannii* on *K. pneumoniae* against cefotaxime in a polymicrobial lung infection. In addition, this protein-mediated protection effect could be coupled to plasmid transfer. The seminal work from Bou and coworkers reported the role of OMVs in transferring the plasmid containing the *bla*_OXA-24/40_ gene between different *A. baumannii* strains (56). Overall, this calls for a deeper understanding of the role of OMVs in polymicrobial infections through studies able to assess the biochemical bases of cross-protection, including the presence of different β-lactamases as selective vesicle cargo.

In closing, this work also underscores the relevance of studying the physiology of β-lactamases in their native bacterial hosts when using model organisms with isogenic backgrounds. The identification of a large, clinically relevant family of β-lactamases as membrane-bound proteins in *A. baumannii* linked to their presence in vesicles requires us to address novel approaches to clinical treatments, particularly in polymicrobial infections.

## MATERIALS AND METHODS

### Bioinformatic analysis

The Entrez module from the Biopython library (56) was used to access the FASTA sequences from the NCBI database using the accession codes listed in the “GenPeptID” column of the β-lactamase database (3). The ignore list command was utilized to exclude entries containing dashes, blank spaces, “assigned”, and “Withdrawn”. For enzymes with multiple accession numbers, the first one listed was selected. All sequences were compiled into a list, which was then saved as a text file. The Python script is available upon request. The generated file was used as the input for the SignalP 6.0 server available at https://services.healthtech.dtu.dk/services/SignalP-6.0. The prediction was executed in slow mode, with “short output” and “Other” selected as the organism option. The results were downloaded as a JSON summary, copied into Supplementary Table 1, and aligned with the information from the β-lactamase database.

### CG-MD simulations

Structures of OXA-23 and OXA-24/40 were obtained from the AlphaFold Protein Structure Database (AFDB) (58) under the UniProt accession codes A0A068J749 and Q8RLA6, respectively (59). This was done as the structures of these enzymes deposited in the Protein Data Bank (PDB) (60) lacked a considerable portion of both proteins N-terminal sequence, including the cysteine residue that is lipidated. Nonetheless, the globular domains of both enzymes’ models were inspected and compared to their deposited structures, confirming that the models had excellent accuracy (OXA-23 AF model pLDDT = 95.7, RMSD against PDB ID 4JF4 = 0.427 Å; OXA-24/40 AF model pLDDT = 95.7, RMSD with PDB ID 3ZNT = 0.208 Å). The models in the AFDB are computed based on the proteins’ complete sequence, so it was necessary to remove the signal peptides of both OXA-23 (up to Cys18) and OXA-24/40 (up to Cys20).

The simulations were performed using the MARTINI 3.0 forcefield (40). The first step involved the coarse-graining of the protein models done using the Martinize 2 tool (61). An elastic network with a bond force constant of 500 kJ mol^−1^ nm^−2^ and lower and upper elastic bond cut-offs of 0.5 and 0.9 nm, respectively, were implemented to stabilize the tertiary structure of the proteins. The parameters and structure of the triacyl moiety covalently attached to the N-terminus cysteine were taken from Rao et al. (62), who also made available a Python script to generate an initial configuration for the lipidated residue. The INSert membrANE (*insane*) program (63) was used to build the simulation systems: a tetragonal box of dimensions 13×13×15 nm^3^ (X, Y, Z) containing a symmetric lipid bilayer of 72% POPE, 16% POPG, and 12 % cardiolipin, mimicking the composition of the *A*. *baumannii* OM inner leaflet (64). The systems were immersed in water with either OXA-23 or OXA-24/40 positioned 7 nm away from the membrane surface. The parameters for cardiolipin were obtained from Corey et al. (65), while the others lipids were already included in the official release of the MARTINI 3.0 force field. The systems were then neutralized with Na^+^ counter ions.

GROMACS (66) version 2022.1 was used to run all the MD simulations. In all stages of the procedure, the cut-off radius for short-range electrostatic and van der Waals interactions was 11 Å; The reaction field method (67) was implemented to calculate long-range electrostatic interactions. Periodic boundary conditions were applied in all directions. Energy minimization and equilibration of the systems was performed as per the recommended protocol of the CHARMM-GUI web server (68) as of July 2023. The production dynamics were performed in the NPT ensemble, with the V-rescale thermostat (69) and the Parrinello-Rahman barostat (70). Both systems had five replicas and each ran for 4 μs. Protein-lipid interaction analyses were performed with the ProLint Web Server (71), using an interaction cut-off of 6 Å. The Visual Molecular Dynamics (VMD) (72) software was used for visual inspection of the simulations and recording of the movies. All protein and membrane images were generated with ChimeraX (73).

### Bacterial strains, culture conditions and plasmid constructions

*Escherichia coli* ATCC 25922 and *Acinetobacter baumannii* ATCC 17978 were used for expression of the empty vector pMBLe-OA and also for expression of the different OXAs. The *bla*_OXA_ genes were cloned into the pMBLe-OA vector (46) fused to a Strep-Tag II sequence and downstream of a pTac promoter inducible by isopropyl β-d-1-thiogalactopyranoside (IPTG). All strains were grown aerobically at 37 °C in lysogeny broth (LB) medium supplemented with gentamicin 20 μg/mL when necessary. Expression of the enzymes were induced at OD = 0.4 by adding 10 µM IPTG and incubated for 4 h at 37 °C.

Chemical reagents were purchased from Sigma-Aldrich.

### Variant constructions by PCR Overlap

The soluble variants of OXA-23 and OXA-24/40 were constructed by site-directed mutagenesis using overlapping primers. For each enzyme we amplified the full length *bla*_OXA_ genes, including their native signal peptides from a pUC57 (Macrogen®) plasmid using mutagenic primers and internal plasmid primers called pMBLe_Fw 5’-GCTGTTGACAATTAATCATCGGCTC-3’ and pMBLe_Rv 5’-CGTAGCGCCGATGGTAGTG-3’.

To construct CA_OXA-23 we used the following couple of primers: pMBLe_Fw 5’-AAATTATGCTGAACCGTAGCACCAGAAAGAAAAAG-3’ and pMBLe_Rv 5’-CTTTTTCTTTCTGGTGCTACGGTTCAGCATAATTT-3’.. The double stranded DNA obtained and the plasmid pMBLe were cleaved using *Nde*I and *Eco*RI and therefore ligated. Then, pMBLe-OXA-23 was digested with *Bam*HI and *Eco*RI and ligated with pMBLe_OA digested with the same enzymes.

To construct CA_OXA-24/40, we used the following series of primers: pMBLe_Fw 5’-AGTTTTAATAGATGAAGCTGCACTGAGAGAAACTAG-3’ and pMBLe_Rv 5’-CTAGTTTCTCTCAGTGCAGCTTCATCTATTAAAACT-3’. The double stranded DNA obtained and the plasmid pMBLe were cleaved using *Nde*I and *Hind*III and were then ligated. Molecular biology enzymes were purchased from Thermo-Fisher®, and primers were ordered from Invitrogen®.

### Cell fractionation

The fractionation protocol was based in the one described by Pettiti et al (74). The cells were pelleted and resuspended in 0.2 M Tris at pH 8, 1M sucrose, 1mM EDTA in a proportion of 10 mL per initial culture liter. Lysozyme 1mg/mL was added and incubated for 5 minutes at room temperature. The suspension was centrifuged during 20 minutes at 30,000 x g, the supernatant content the periplasmic fraction and the pellet the spheroplast and membrane remains. The pellet was resuspended in 10mM Tris at pH 8.5 EDTA 2.5 mM 10 μg/mL DNAse 1mM PMSF and sonicated. Once cell debris were removed, the suspension was ultracentrifuged at 45,000 x g for 45 minutes. The supernatant containing the cytoplasm was collected. The pelleted total membrane was resuspended in 10 mM Hepes, 0.2 M NaCl at pH 7.5. The resuspension volumes were normalized according to the OD_600_ and the initial culture volume.

### Selective membrane protein solubilization

Membrane proteins were extracted sequentially. Firstly, the total membrane fraction was pelleted by ultracentrifugation at 45,000 x g during 45 min and gently resuspended in cold 1 M KCl. After 30 min incubation on ice they were ultracentrifuged again. The supernatants containing loosely associated peripheral proteins were collected and the pellets were resuspended in 0.1 M Na_2_HCO_3_ at pH 11.5. After a 30min incubation on ice to release peripheral proteins associated by strong electrostatic interactions were centrifugated again. The pellets were resuspended in 1% w/v Triton X-100 and incubated 30 min on ice to extract integral or hydrophobically-associated proteins in detergent micelles. The supernatants were collected, and the pellets were resuspended in 10 mM Hepes, 0.2 M NaCl at pH 7.5.

### Protein Immunodetection

Immunoblotting assays were realized in PVDF membranes using Strep-Tag® II monoclonal antibodies (at 1:1000 dilution from a 200 μg/ml solution) (Novagen®) and immunoglobulin G-alkaline phosphatase conjugates (at 1:5000 dilution). Monoclonal anti-RNApol was used as a cytoplasmic control and polyclonal anti-OmpA kindly provided by Dr. Alejandro Viale was used as a membrane marker.

The whole cells, periplasm and cytoplasm samples were normalized according to the following equation: V = 100 μL x OD_600_ x V_c_, where V_c_ is the starting volume of culture sample. Total membranes fraction and OMVs were normalized according to a lipid quantification by FM4-64 (Thermofisher®). Protein band intensities were quantified by using Gel Analyzer software (75).

### Purification of OMVs and OXAs levels detection into OMVs

Overnight cultures of *A. baumannii* pMBLe-OA-*bla* cells were grown in 300 mL of LB broth at 37°C, reaching an OD_600_ of 0.4. Subsequent induction with 20 μM IPTG was followed by continued overnight growth with agitation. The cells were harvested, and the supernatant was filtered through a 0.45-μM membrane (Millipore). Ammonium sulfate was added to the filtrate at a concentration of 55% (w/vol), followed by overnight incubation with stirring at 4 °C. Precipitated material was separated by centrifugation at 12,800 × g for 10 min, resuspended in 10 mM HEPES, 200 mM NaCl at pH 7.4, and dialyzed overnight against >100 volumes of the same buffer. Next, samples were filtered through a 0.45-μM membrane, layered over an equal volume of 35% (w/vol) sucrose solution, and ultracentrifuged at 150,000 × g for 1 h and 4 °C. Pellets containing the OMVs were washed once with 10 mM HEPES, 200 mM NaCl at pH 7.4, and stored at −80 °C until use.

The OMVs were quantified by two different methods. The total protein concentration was measured by the Pierce bicinchoninic acid (BCA) protein assay kit (Thermo Scientific®) as described (37). Lipid content associated with OMVs was determined using the lipophilic fluorescent dye FM4-64 (Thermofisher®) as described previously (76). Briefly, a portion of OMVs was incubated with FM4-64 (2 μg/ml in PBS) for 10 min at room temperature. Separate samples of OMVs and the FM4-64 probe were used as negative controls. After excitation at 515 nm, emission at 640 nm was measured with the multiplate reader SYNERGY HT (Biotek®).

The OMV content was analyzed by SDS-PAGE and immunoblotting. Gel lanes were equally loaded based on total protein and lipid content. The pre-stained *Blue Plus® II* Protein Marker (14-120 kDa) provided molecular weight standards for Fig. 4. To determine the levels of OXA-23, CA_OXA-23, OXA-24/40, and CA_OXA-24/40 in OMVs, the mature protein band intensities in whole cells (WC) and in the OMVs, derived from *A. baumannii* expressing each OXA protein, were quantified from polyvinylidene difluoride (PVDF) membranes using GelAnalyzer software (53). The quantity of each OXA protein in the OMVs (from immunoblots) was divided by the quantity of each OXA in whole cells (from immunoblots). Finally, the values plotted in Fig. 4B (expressed as a percentage) correspond to the normalization of the levels of each soluble OXA (CA_OXA-23 or CA_OXA-24/40) to the value of its corresponding wild-type OXA (OXA-23 or OXA-24/40), which was taken as 100 percent.

### Proteinase K accessibility assay

OMVs were resuspended in a buffer containing 10 mM Tris HCl (pH 8) and 5 mM CaCl_2_. When required, OMVs were lysed by incubation with 1 % (vol/vol) Triton X-100 for 30 min at 37 °C. Intact and lysed OMVs were incubated for 60 min at 37 °C in the presence of 100 μg/ml proteinase K. The reaction was stopped by the addition of 5 mM phenylmethanesulfonyl fluoride (PMSF), and samples analyzed by SDS-PAGE and immunoblotting.

### MICs of OXA-23, CA_OXA-23, OXA-24 and CA_OXA-24

To determine minimum inhibitory concentrations (MIC, µg/ml) of the β-lactam antibiotics imipenem and piperacillin (Sigma-Aldrich®) on strains of *A. baumannii* carrying OXAs we followed the standard agar plate protocol recommended by the CLSI.

### Effect of OMVs loaded with OXAs on minimum inhibitory concentrations (MIC) of bacteria

To determine the minimum inhibitory concentrations (MIC, µg/ml) of the β-lactam antibiotics imipenem and piperacillin (Sigma-Aldrich®) on β-lactam-susceptible strains of *A. baumannii*, *E. coli*, and *P. aeruginosa* after treatment with OMVs, we utilized OMVs purified from *A. baumannii* carrying an empty vector (EV) or expressing either lipidated (OXA-23 or OXA-24/40) or soluble enzymes (CA_OXA-23 or CA_OXA-24/40). We determined the MIC by broth-dilution method in 96-well plates according to the CLSI guidelines. β-lactam-susceptible cells (5 × 10^5^ CFU/ml) were inoculated into medium without β-lactam or with 2-fold increasing concentrations of imipenem or piperacillin, and with 1 μg/ml of OMVs from *A. baumannii* carrying an empty vector (EV) or expressing OXAs. The 96-well plates were incubated at 37 °C with constant shaking and the OD was recorded at 600 nm at 30 min time intervals, using a Biotek Epoch 2 microplate reader. MIC values were measured from two independent experiments.

## ACKNOWLEDGMENTS

This research was supported by grants from the National Institutes of Health (R01AI100560 to R.A.B. and A.J.V.), Agencia I+D+I (PICT-2020-00031 to A.J.V.), MinCyT (REPARA to A.J.V.), the European Union’s Horizon 2020 research and innovation programme under the Marie Skłodowska-Curie (grant agreement No. 945363 to F.A.M.), and the Swiss National Science Foundation (CRSII5_198737 to M.D.P.). A.J.V. and C.L. are staff members from CONICET. L.C. is recipient of fellowship from CONICET, Argentina. F.A.M. and M.D.P. are staff members of EPFL. We thank Marina Avecilla (IBR-CONICET) for her excellent technical assistance.

C.L. and A.J.V. designed research and supervised the study. L.C. and G.B. performed the bioinformatics analysis of the lipobox sequences. L.C. performed plasmid constructions for OXAs and their soluble variants, cell fractionation and protein localization, selective membrane solubilization and OXA immunodetection. C.L. performed OMVs purification, detection and determination of OXA levels into OMVs, proteinase K assay and protection assay with OMVs. F.A.M. performed the molecular simulation experiments. L.C., C.L. and F.A.M. designed the figures. All authors analyzed data, discussed results and contributed and edited manuscript. The content is solely the responsibility of the authors and does not necessarily represent the official views of the National Institutes of Health or the Department of Veterans Affairs.

